# Mechano-regulation of bone adaptation is controlled by the local *in vivo* environment and logarithmically dependent on loading frequency

**DOI:** 10.1101/2020.05.15.097998

**Authors:** Ariane C. Scheuren, Paul Vallaster, Gisela A. Kuhn, Graeme R. Paul, Angad Malhotra, Yoshitaka Kameo, Ralph Müller

## Abstract

It is well established that cyclic, but not static, mechanical loading has anabolic effects on bone. However, the function describing the relationship between the loading frequency and the amount of bone adaptation remains unclear. Using a combined experimental and computational approach, this study aimed to investigate whether bone mechano-regulation is controlled by mechanical signals in the local *in vivo* environment and dependent on loading frequency. Specifically, by combining *in vivo* micro-computed tomography (micro-CT) imaging with micro-finite element (micro-FE) analysis, we monitored the changes in microstructural as well as the mechanical *in vivo* environment (strain energy density (SED) and SED gradient) of mouse caudal vertebrae over 4 weeks of either cyclic loading at varying frequencies of 2Hz, 5Hz, or 10Hz, respectively or static loading. Higher values of SED and SED gradient on the local tissue level led to an increased probability of bone formation and a decreased probability of bone resorption. In all loading groups, the SED gradient was superior in the determination of local bone formation and resorption events as compared to SED. Cyclic loading induced positive net remodeling rates when compared to sham and static loading, mainly due to an increase in mineralizing surface and a decrease in eroded surface. Consequently, bone volume fraction increased over time in 2Hz, 5Hz and 10Hz (+15%, +21% and +24%, p<0.0001), while static loading led to a decrease in bone volume fraction (−9%, p≤0.001). Furthermore, regression analysis revealed a logarithmic relationship between loading frequency and the net change in bone volume fraction over the four week observation period (R^2^=0.74). In conclusion, these results suggest that bone adaptation is regulated by mechanical signals in the local *in vivo* environment and furthermore, that mechano-regulation is logarithmically dependent on loading frequency with frequencies below a certain threshold having catabolic effects, and those above anabolic effects. This study thereby provides valuable insights towards a better understanding of the mechanical signals influencing bone formation and resorption in the local *in vivo* environment.

## Introduction

It is well established that cyclic, but not static loading has anabolic effects on bone [1–4]. This clear-cut discrepancy in osteogenic responses to both loading patterns highlights the key role of loading frequency in mechano-regulation of bone remodeling - the coordinated process by which bone is continuously formed and resorbed. Yet, the exact relationship between loading frequency and bone remodeling and bone adaptation remains unclear. While both experimental [5–7] and theoretical studies [8, 9] have suggested a dose-response relationship such that bone formation increases with higher loading frequencies, Warden and Turner have shown this relationship to be non-linear [10] using an axial loading model of mouse ulnae. Using this model, they showed that cortical bone adaptation increased with frequencies up to 5 and 10Hz, but then plateaued thereafter. In line with these results, more recent *in silico* studies have found non-linear relationships between loading frequency and bone adaptation both in cortical [11] as well as in trabecular [12] bone. In the latter study, a single trabecula was subjected to cyclic uniaxial loading at frequencies of either 1Hz, 3Hz, 5Hz, 10Hz or 20Hz. Similar to the study by Warden et al., bone volume fraction increased up to 10Hz but then plateaued thereafter [12]. However, owing to the lack of *in vivo* studies investigating the effects of loading frequency on trabecular bone adaptation, the validity of such *in silico* studies remains unclear. Furthermore, as frequency effects have been shown to vary depending on the anatomical region investigated [13], the optimal frequency must be identified for every specific loading model.

Using a tail-loading model, we have previously shown that cyclic loading at a frequency of 10Hz over four weeks elicits anabolic responses in mouse caudal vertebrae [14]. Furthermore, by combining time-lapsed micro-computed tomography (micro-CT) imaging with micro-finite element (micro-FE) analysis, we were able to demonstrate that bone remodeling in the trabecular compartment is controlled by local mechanical signals at the tissue level [15–17]. Specifically, by registering consecutive time-lapsed *in vivo* micro-CT images onto one another [18], sites of bone formation and resorption were quantified in three dimensions and subsequently linked to corresponding mechanical signals calculated in the local *in vivo* environment (L*iv*E) [15–17]. Herein, simulating the distribution of strain energy density (SED) - defined as the increase in energy associated with the tissue deformation per unit volume (i.e., a measure of direct cell strain) - within the caudal vertebrae revealed that bone formation was more likely to occur at sites of high SED, whereas bone resorption was more likely to occur at sites of low SED [15, 17]. While SED is widely used as a mathematical term to describe the mechanical signal influencing bone remodeling [15, 19–21], other mechanical signals, such as interstitial fluid flow through the lacuna-canalicular network (LCN), are also known to play a major role in determining the local mechanical environment surrounding osteocytes, the main mechanosensors in bone [22–24]. In this respect, it has been suggested that measures of fluid flow, such as the gradient in SED, would allow improved predictions of adaptive bone remodeling events [16, 25]. In this study, we therefore aimed to 1) investigate the effects of varying loading frequencies on the mechano-regulation of trabecular bone in mouse caudal vertebrae, 2) assess whether adaptive bone remodeling can be linked to mechanical signals in the local *in vivo* environment and 3) compare the modeling performance of SED and the gradient in SED for the prediction of local bone formation and resorption events on the tissue level. Specifically, we used time-lapsed *in vivo* micro-CT imaging to monitor bone adaptation over time in individual animals in response to cyclic loading at frequencies of 2Hz, 5Hz and 10Hz as well as in response to static loading. In comparison to conventional two-dimensional (2D) histomorphometric techniques, which have previously been used to investigate effects of varying frequencies on bone adaptation [1, 4, 7, 13], the ability to quantify not only bone formation but also resorption over time could elucidate contrasting effects observed after static and cyclic loading. Furthermore, the analysis of various mechanical signals in the local *in vivo* environment by means of micro-FE analysis provided a better understanding of these signals influencing bone forming and resorbing cells on the local level. Finally, by determining the conditional probabilities for bone formation and resorption events to occur as a function of these mechanical signals [15], this study contributed towards the description of the relationship between local mechanical signals and the subsequent mechano-regulation of bone adaptation. In future, these results will be highly beneficial for *in silico* studies aiming to predict the mechano-regulation of bone adaptation in response to various interventions.

## Results

### Bone adaptation to load is dependent on loading frequency

In order to investigate the effects of varying loading frequencies on bone adaptation, we used an *in vivo* micro-CT approach [26] to monitor bone adaptation of the sixth caudal vertebrae of C57BL/6J mice subjected to a 4-week loading regime of either sham (0N), 8N static or 8N cyclic loading with frequencies of 2Hz, 5Hz, or 10Hz, respectively. Table 1 shows the difference between the first and last time point (i.e., bone parameter_week4-week0_) of the bone structural parameters in the trabecular and cortical bone. In the trabecular bone compartment, the difference of bone volume fraction (BV/TV) and trabecular thickness (Tb.Th) between the first and last time point was significantly different between groups (p<0.0001), whereas no significant differences were detected between groups for the trabecular number and separation (Tb.N and Tb.Sp, p>0.05). Whereas the sham and static loading groups showed a net decrease in BV/TV and Tb.Th, the cyclic loading groups at 2Hz, 5Hz and 10Hz displayed increases in BV/TV and Tb.Th, with all of them being significantly different to the sham group (Table 1). With respect to the structural parameters of cortical bone, differences between the first and last time point were significantly different between groups for cortical area fraction (Ct.Ar/Tt.Ar, p<0.0001) and cortical thickness (Ct.Th, p<0.01), where the cyclic loading groups showed significantly greater increases compared to the sham-loaded group (Table 1).

**Table 1.** Difference between week 0 and week 4 for bone structural parameters in the trabecular and cortical compartments. P-values denote a significant difference between groups determined by one-way ANOVA, while “*” denotes significant difference to sham as assessed by multiple comparisons Dunnett’s test (* p<0.05, ** p<0.01, *** p<0.001 and **** p <0.0001).

| Morphometric parameter | sham | static | 2Hz | 5Hz | 10Hz | p-value |
| --- | --- | --- | --- | --- | --- | --- |
| **BV/TV (%)** | -0.93±0.789 | -1.408±1.392 | 2.333±1.315**** | 3.240±1.692**** | 3.680±1.084**** | <0.0001 |
| **Tb.Th (mm)** | 0.006±0.005 | 0.006±0.005 | 0.021±0.01*** | 0.020±0.007** | 0.021±0.006*** | <0.0001 |
| **Tb.N (1/mm)** | 0.263±0.122 | -0.286±0.076 | -0.234±0.127 | -0.341±0.082 | -0.295±0.221 | >0.05 |
| **Tb.Sp (mm)** | 0.034±0.019 | 0.037±0.009 | 0.031±0.024 | 0.043±0.013 | 0.034±0.027 | >0.05 |
| **Ct.Ar/Tt.Ar (%)** | 0.507±1.187 | 0.524±1.931 | 2.746±0.950* | 3.838±2.209** | 3.496±1.733** | <0.0001 |
| **Ct.Th (mm)** | 0.004±0.005 | 0.005±0.005 | 0.013±0.005* | 0.014±0.013* | 0.014±0.009* | <0.01 |

Figure 1 shows the relative changes in trabecular bone morphometric parameters over the 4-week loading period for the different loading groups. BV/TV developed differently over time between the loading groups (interaction effect, p<0.0001). Compared to the sham-loaded group, which showed no change in BV/TV over time (−6%, p>0.05), cyclic loading at all frequencies (2Hz, 5Hz and 10Hz) led to a dose-response increase in BV/TV with higher frequencies resulting in higher increases in BV/TV (Fig 1A). Herein, the 5Hz and 10Hz groups showed a significant increase compared to baseline already 2 weeks after the start of loading (p≤0.001 and p<0.0001), while the 2Hz group showed a significant increase relative to baseline only after three weeks (p≤0.001). At the end of the 4-week loading regime, these groups showed a 15%, 21% and 24% higher BV/TV relative to baseline (p<0.0001 for 2Hz, 5Hz and 10Hz). Static loading on the other hand, had catabolic effects resulting in significantly lower BV/TV (−9%, p≤0.01) at the last time point relative to baseline. In line with the changes in BV/TV, Tb.Th developed differently over time between the loading groups (interaction effect, p<0.0001, Fig 1B). By the end of the 4-week loading intervention, all cyclic loading groups showed significant increases in Tb.Th (p<0.0001), which was not observed in the static and sham-loaded groups (p>0.05). Although the number of trabeculae (Tb.N) decreased and trabecular separation increased (Tb.Sp) over time (Fig 1C,D, p<0.001), no relative differences were observed between the groups (p>0.05). These results thus suggest that increases in BV/TV due to cyclic loading were mainly driven by thickening of the trabeculae rather than by the inhibition of the reduction in the number of trabeculae.

**Fig 1.**
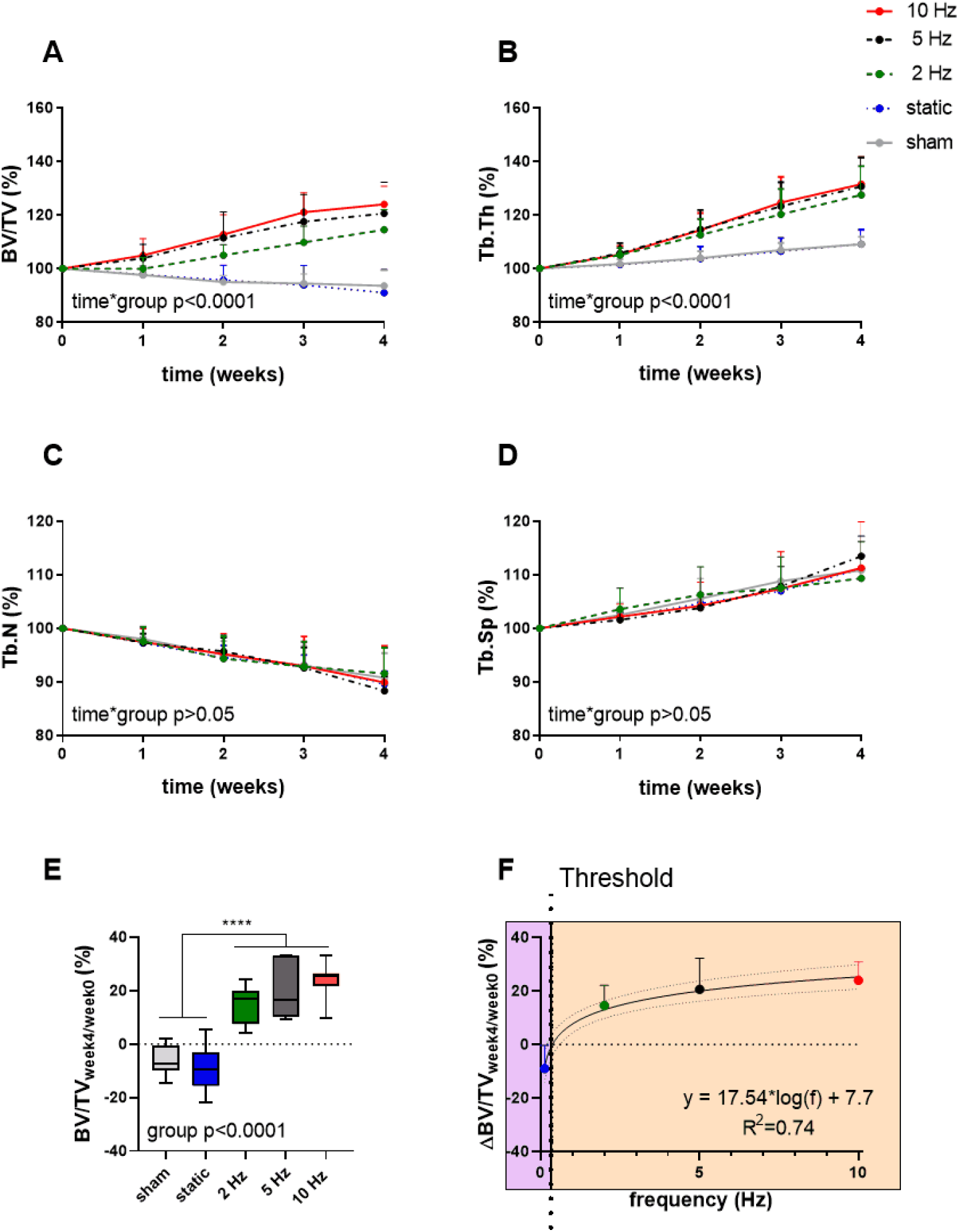
Relative changes of structural bone morphometric parameters in the trabecular compartment over the 4-week loading period as assessed by *in vivo* micro-CT. (A) Bone volume fraction (BV/TV), (B) trabecular thickness (Tb.Th), (C) trabecular number (Tb.N) and (D) trabecular spacing (Tb.Sp). (Data represent mean±standard deviation (SD) for n=5-8/group, p-values for interaction effect between group and time are shown as determined by linear mixed effects model). (E) The relative change from week 4 relative to baseline (BV/TV_week4/week0_) (F) was fitted with a logarithmic regression line. (Data represent mean±SD for n=5-8/group, p-value for main effect of group determined by one-way ANOVA, **** p<0.0001 denotes significant difference between groups determined by post hoc Tukey’s multiple comparisons test).

By plotting the relative changes in BV/TV as a function of loading frequency, regression analysis revealed a logarithmic relationship between bone adaptation and loading frequency as a best fit to the data (R^2^=0.74, Fig 1F) with loading frequencies above 0.36Hz±0.08 having anabolic effects, and frequencies below this threshold having catabolic effects. Although there were no significant differences between the cyclic loading groups, loading at 10Hz had the earliest and largest anabolic effects compared to the other frequencies.

Aside from providing information on changes in bone structural parameters over time, *in vivo* micro-CT also provided the possibility to assess dynamic bone formation and resorption activities such as bone formation/resorption rate (BFR/BRR), mineral apposition/resorption rate (MAR/MRR) and mineralizing/eroded surface (MS/ES) [18]. The net remodeling rate (BFR-BRR), which gives an indication whether there was overall bone gain (i.e., BFR-BRR>0) or loss (i.e., BFR-BRR<0) occurring within the trabecular compartment, tended to develop differently between groups (p=0.056). Compared to the static and sham-loaded groups, which had an overall negative remodeling balance, the 2Hz, 5Hz and 10Hz had an overall positive remodeling balance (p≤0.01, p≤0.001 and p<0.0001, Fig 2A). The net remodeling rate did not significantly change over time. When bone formation and resorption rates were analyzed separately, the main differences in the cyclic loading groups were in the reduced BRR as compared to the sham and static groups. While BFR did not significantly differ between groups (p>0.05, Fig 2B), BRR was 35% (p<0.01), 50% (p<0.0001) and 44% (p<0.0001) lower in the 2Hz, 5Hz, 10Hz groups, respectively, compared to the sham-loaded group (Fig 2C). The static group on the other hand had a similar BRR (−2%, p>0.05) as the sham-loaded group (Fig 2C).

**Fig 2.**
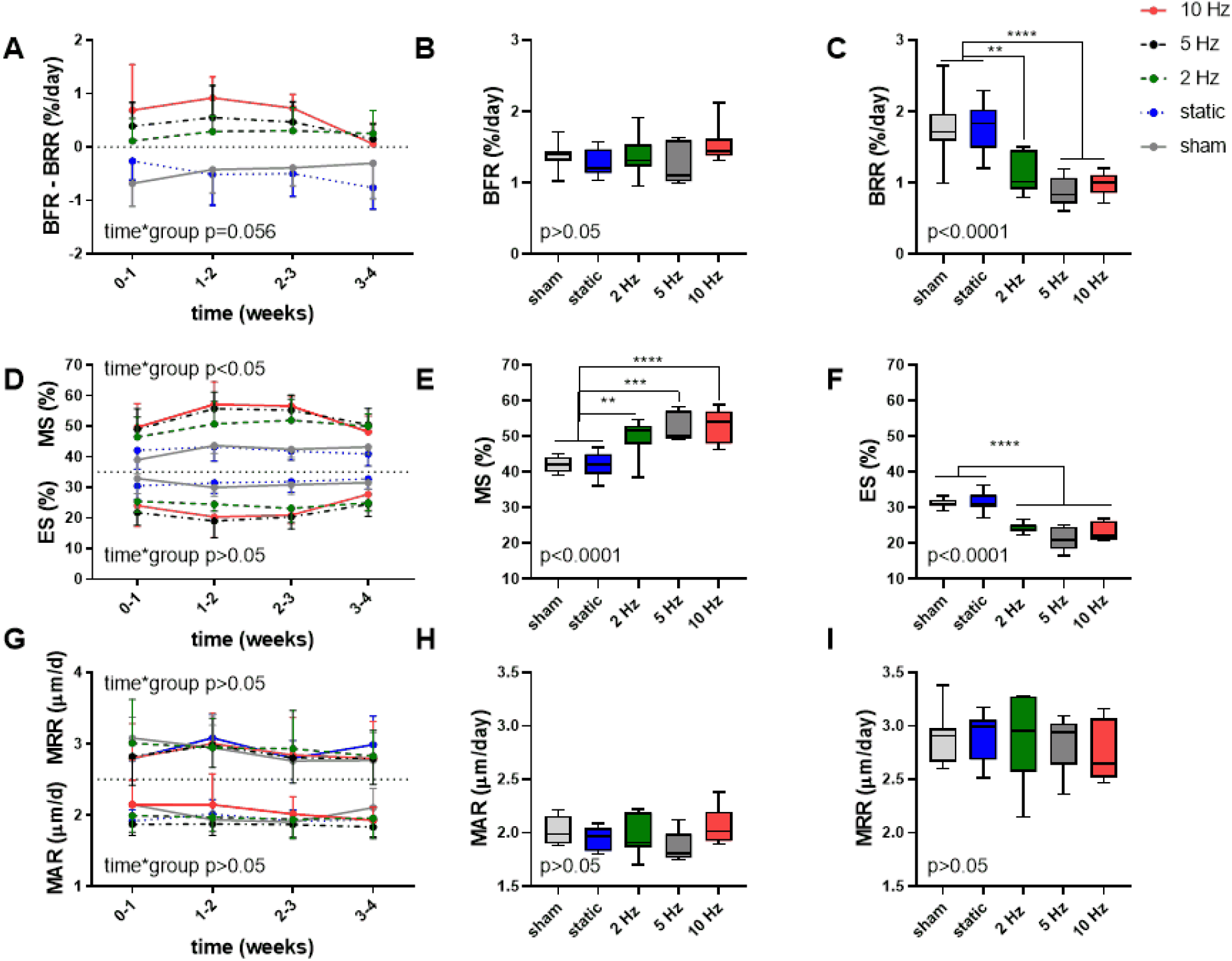
Dynamic bone morphometric parameters in the trabecular compartment in the different loading groups as assessed by *in vivo* micro-CT. (A) Changes in the net remodeling rate shown as the difference between bone formation rate (BFR) and bone resorption rate (BRR) over the 4-week loading period. Overall difference between groups of (B) BFR and (C) BRR. (D) Mineralized surface (MS) and eroded surface (ES) over the 4-week loading period. Overall difference between groups of (E) MS and (F) ES. (G) Mineral apposition rate (MAR) and mineral resorption rate (MRR) over the 4-week loading period. Overall difference between groups of (H) MAR and (I) MRR. (Data represent mean±SD for n=5-8/group, p-values for interaction effect between group and time are shown as determined by linear mixed effects model (A,D,G), boxplots showing the differences between groups as determined by Tukey’s post hoc multiple comparisons test * p<0.05, ** p<0.01, *** p<0.001, **** p<0.0001 (B,C,E,F,H,I))

A difference between the cyclic and static loading groups was also apparent when investigating the surfaces of formation (mineralized surface, MS, interaction effect p<0.05) and resorption (eroded surface, ES, interaction effect p≥0.05) sites with the cyclic loading groups having a higher MS and lower ES compared to the static and sham-loaded groups (Fig 2D). On average, formation sites occupied 2, 2.5 and 2.6 more surfaces than resorption sites for the 2Hz, 5Hz and 10Hz groups, and only 1.4 times more for the control and static groups, respectively.

Furthermore, the 2Hz, 5Hz and 10Hz groups had a 18% (p=0.0078), 25% (p=0.0007) and 26% (p>0.0001) higher mineralized surface (MS) and a 22% (p<0.0001), 32% (p<0.0001) and 26% (p<0.0001) lower eroded surface (ES) compared to the sham-loaded group, while the static group had similar MS and ES compared to sham-loading (p>0.05, Fig 2E-F). The mineral apposition and resorption rates (MAR and MRR), which represent the thicknesses of formation and resorption packages, respectively, did not develop differently between groups (interaction effects p=0.586 and p=0.459). Furthermore, the MAR and MRR were similar between groups (p>0.05), thus suggesting that they are not affected by loading (Fig 2G-I). This indicates that cyclic loading had a greater effect on surface than on thickness of formation as well as resorption sites.

### Bone adaptation to load is controlled by mechanical signals in the local *in vivo* environment

In order to assess whether bone remodeling events - namely formation, quiescence (i.e., where no remodeling occurred) and resorption - can be linked to the corresponding mechanical signals in the local *in vivo* environment, we performed micro-finite element (micro-FE) analysis to calculate the strain distribution within the tissue. As deformation (direct cell strain) and interstitial fluid flow (shear stress) are hypothesized to be the main mechanical stimuli that regulate load-induced bone adaptation [27], we quantified the strain energy density (SED) magnitudes as a measure of mechanical deformation and the spatial gradient thereof (∇SED), as a measure of fluid flow, respectively [16, 28]. Figure 3 displays a representative visualization of a section of the vertebrae of the 10Hz group showing sites of bone remodeling (Fig 3A) as well as the corresponding maps of SED (Fig 3B) and ∇SED (Fig 3C). From this qualitative analysis, it is apparent that bone resorption occurs at sites of lower SED and ∇SED, respectively, whereas bone formation occurs at sites of higher SED and ∇SED (Fig 3).

**Fig 3.**
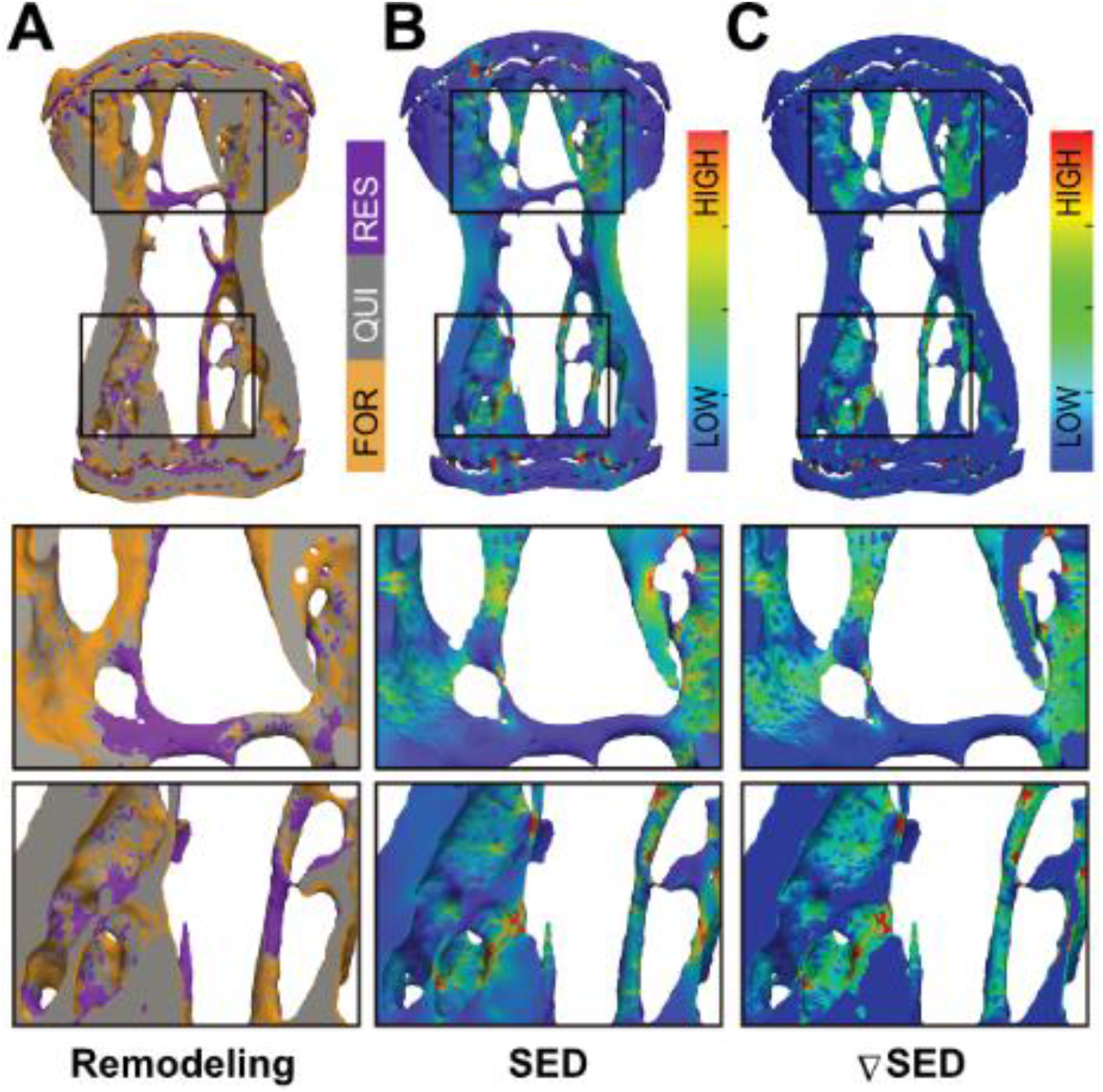
Qualitative visualization linking bone remodeling sites (formation, quiescence, resorption) with the mechanical environments *in vivo*. (A) Overlay of time-lapsed micro-CT images showing sites of bone formation (orange), quiescence (grey) and resorption (purple). Corresponding map of the (B) strain energy density (SED) and (C) gradient thereof (∇SED) showing sites of higher (red) and lower (blue) SED/∇SED values obtained by micro-finite element (micro-FE) analysis.

To establish a quantitative description of the mechano-regulation of bone remodeling, we calculated the conditional probabilities for a given remodeling event to occur as a function of the mechanical stimuli, also known as remodeling rules [15]. Figure 4 shows the conditional probability curves for formation (orange), quiescence (grey) or resorption (purple) to occur at a given value of SED (Fig 4A,C,E) or ∇SED (Fig 4B,D,F) for the different groups averaged over all time points. For all groups, the conditional probability for bone formation to occur was higher at higher values of SED and ∇SED, respectively (SED/SED_max_ > 0.18) whereas bone resorption was more likely to occur at lower values (SED/SED_max_ < 0.18). The probability curves for all groups were fit by exponential functions (Table S1), of which the coefficients provide information on the functioning of the mechanosensory system as described previously [15]. When comparing the slopes of the formation probability curves (parameter a, Fig 4A,B and Table S1), which can be interpreted as the mechanical sensitivity of the system, there was a gradual increase of the mechanical sensitivity with increasing frequency with the 10Hz group showing the highest mechanical sensitivity (a_SED_ = 0.217, a_SEDgrad_ = 0.316). For the resorption probability curves (Fig 4E,F and Table S1), the 5Hz and 10Hz groups showed similar mechanical sensitivity to SED (a_SED_ = 0.284), while the 5Hz group showed highest sensitivity to ∇SED (a_SEDgrad_ = 0.264 compared to a_SEDgrad_ = 0.252 in 10Hz group). The probability of the quiescence however, was not influenced by loading frequency (Fig 4C,D). When comparing between SED and ∇SED as mechanical stimuli driving bone remodeling events, it seems that in all groups, formation was more sensitive to ∇SED shown by the higher slopes (a_SED_ < a_gradSED_) of the probability curves (Fig 4A,B and Table S1). In contrast, resorption seemed to be more sensitive to SED (a_SED_ > a_gradSED_, Fig 4 E,F and Table S1).

**Fig 4.**
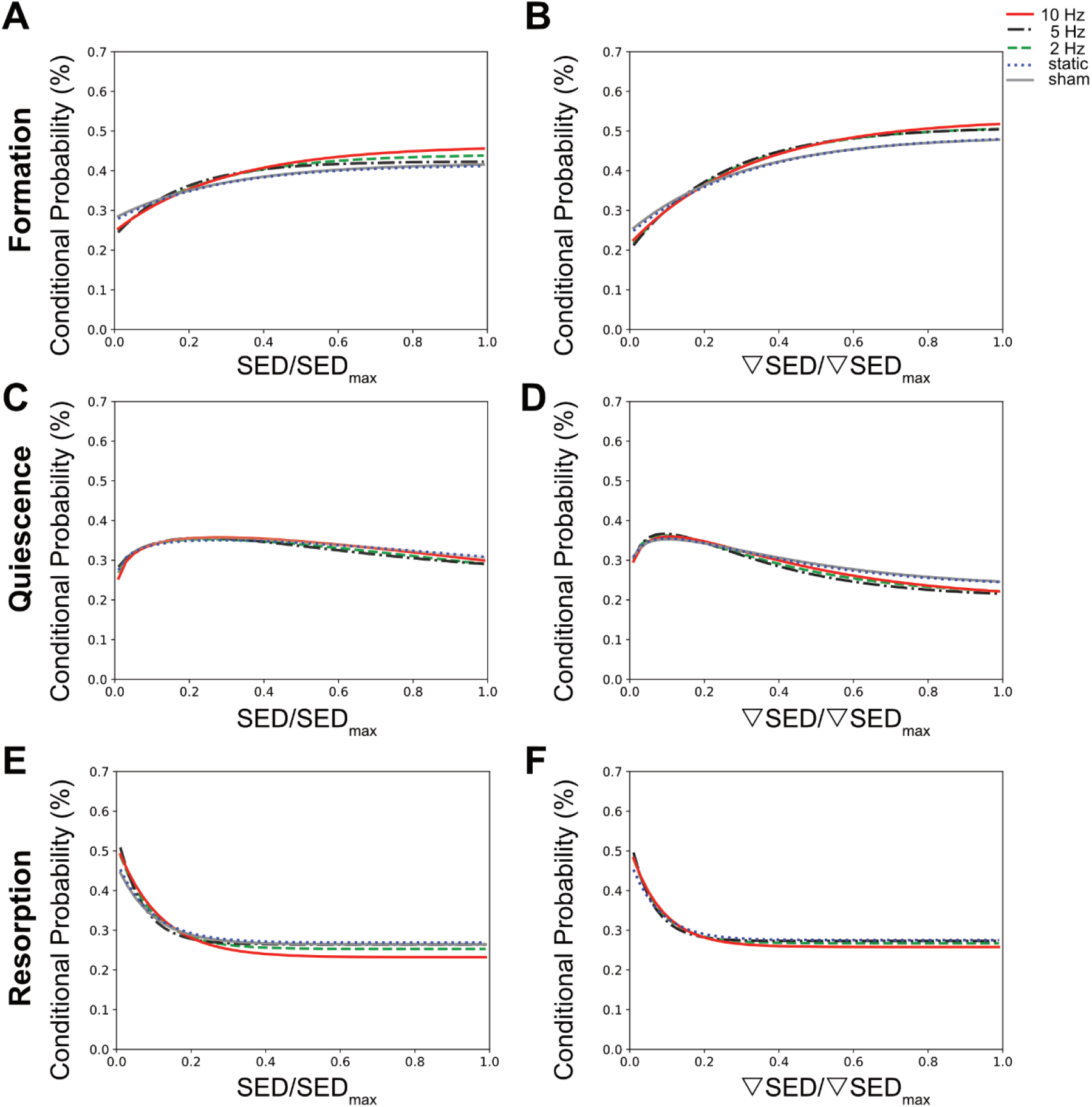
Conditional probabilities connecting SED (left side) and SED gradient (∇SED, right side) with remodeling events. The plots show the exponential fitting functions for (A,B) bone formation (top row), (C,D) quiescence (middle row) and (E,F) resorption (bottom row) in all the loading groups averaged over all time points.

To better compare the modeling performance of SED versus ∇SED for the prediction of bone remodeling events, an area under the receiver operator characteristic curve (AUC) approach was used (Fig 5). For all groups, the AUC values for formation (for all groups p<0.0001, Fig 5A) and resorption (for all groups p<0.05 except for 5Hz p<0.10, Fig 6C) events were higher for the ∇SED compared to SED. No difference between SED and ∇SED was observed for quiescence (Fig 5B). These results suggest that ∇SED has a better modeling performance compared to SED for determining the probability of bone formation and resorption events.

**Fig 5.**
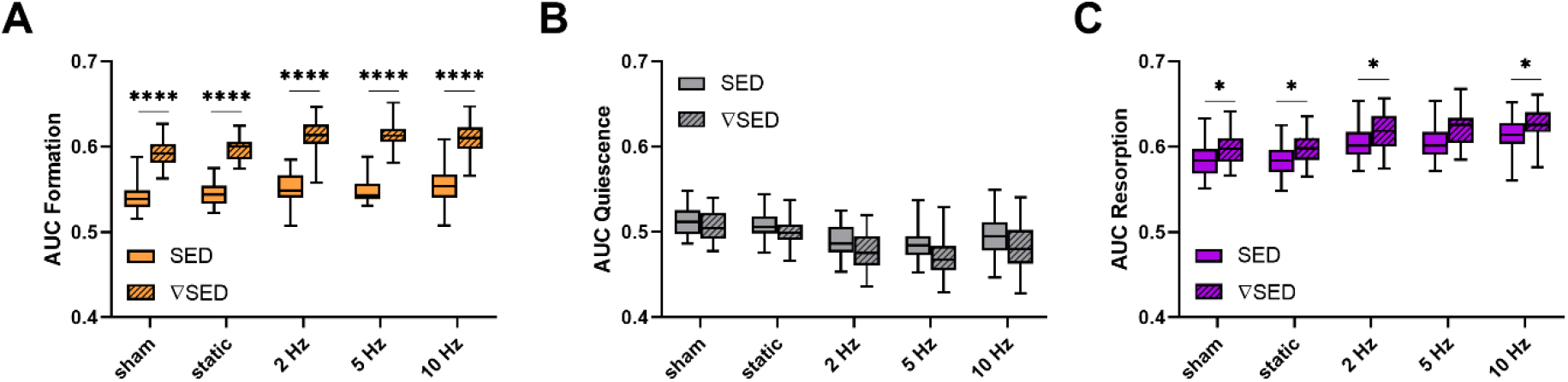
Area under the curve (AUC) values for the comparison of the modeling performance of SED and SED gradient. (A) Formation (orange), (B) quiescence (grey) and (C) resorption (violet) sites for the different loading groups comparing modeling performance of SED (solid bars) and SED gradient (∇SED, striped bars). (Boxplots for n=5-8/group, * p<0.05, **** p<0.0001 differences between groups determined by Tukey’s multiple comparisons test).

## Discussion

In this study, the effects of cyclic loading at varying frequencies as well as of static loading on trabecular bone adaptation in mouse caudal vertebrae were investigated. Furthermore, using a combination of *in vivo* micro-CT and micro-FE analysis, we assessed whether local bone remodeling events (formation and resorption) can be linked to diverse mechanical environments *in vivo*.

While static loading had catabolic effects, cyclic loading at 2Hz, 5Hz and 10Hz had anabolic effects on trabecular bone. In line with previous studies using the tail loading model [17, 26], cyclic loading over four weeks led to an increase in BV/TV, which was driven by the thickening of individual trabeculae rather than a prevention of loss in trabecular number. Furthermore, by registering consecutive time-lapsed images onto one-another, we were able to quantify both bone formation as well as bone resorption activities in three dimensions [26], which to the best of our knowledge, has not yet been used to assess the effects of static loading regimes. Specifically, we showed that cyclic loading mainly affects the surfaces of the bone formation and resorption sites (MS and ES), rather than the thickness of these remodeling packets (MAR and MRR). In agreement with previous studies [18, 26], these results suggest that cyclic loading promotes osteoblast recruitment, while simultaneously inhibiting osteoclast recruitment. Ultimately, cyclic loading results in larger mineralized surfaces and smaller eroded surfaces while keeping the thickness of the remodeling packets constant.

Notably, this study showed a logarithmic relationship between loading frequency and load-induced bone adaptation with frequencies above a certain threshold having anabolic effects and those below having catabolic effects. That cyclic, but not static loading, has anabolic effects on cortical bone has been shown in various animal models including rabbits [2], turkeys [1] and rats [3, 4]. However, to the best of our knowledge, the effect of static loading has not yet been assessed in trabecular bone in mice. In line with the existence of a frequency threshold (0.36Hz±0.08) to elicit anabolic responses as demonstrated in this study, Turner et al. found that bone formation rate in rat tibiae only increased with frequencies above 0.5Hz, followed by a dose-response increase up to 2Hz [5]. Using a similar design as our study, Warden et al. showed increased cortical bone adaptation with increasing loading frequencies up to 5 to 10Hz with no additional benefits beyond 10Hz [10]. In a theoretical study, Kameo et al. furthermore showed similar results by subjecting individual trabeculae to uniaxial loading at frequencies ranging from 1 to 20Hz [12]. Although one would expect higher loading frequencies to lead to higher cellular stimulation and a consequent greater anabolic response, it has been suggested that frequencies above a certain threshold (10Hz) reduce the efficiency of fluid flow through the LCN, thus resulting in inefficient mechanotransduction [10, 29]. More recently, by monitoring Ca^2+^ signaling in living animals, Lewis et al. have shown that osteocyte recruitment was strongly influenced by loading frequency [30]. Another physiological system, for which the relationship between frequency and mechanotransduction is widely studied, is the inner ear [31, 32]. Hair cells, the cells responsible for transducing mechanical forces originating from acoustic waves to neural signals, are sensitive to frequency [31, 33]. Furthermore, the sensitivity of the ear varies with the frequency of sound waves resulting in a limited range of frequencies that can be perceived. Hence, drawing an analogy to the theory of sound pressure level, which also displays logarithmic laws [34], it is possible that bone’s response to frequency is similar to the perception of sound in human hearing.

One limitation of this study was that loading at low (1Hz) and higher (>10Hz) frequencies was not assessed. Furthermore, as the strain magnitude and duration of individual loading bouts were the same for all loading groups, the number of cycles and strain rate differed between the different loading groups. From this study design, it therefore remains impossible to know whether the number of cycles or the loading frequency are the main factors driving load-induced bone adaptation. Hence, whether bone’s osteogenic response to loading is indeed limited to a specific range of frequencies, below and above which bone becomes less osteogenic, requires further *in vivo* experiments.

Using the combined approach of time-lapsed *in vivo* micro-CT imaging and micro-FE analysis, we showed that bone remodeling activities were correlated to the local mechanical environment at the tissue level. In agreement with previous studies [15, 17], bone formation was more likely to occur at sites of higher SED whereas bone resorption was more likely to occur at sites of lower SED. Furthermore, compared to static loading, cyclic loading decreased the probability of non-targeted bone remodeling, which led to an increase in bone formation and a decrease in bone resorption. In addition, we showed that the SED gradient was better at predicting bone formation and resorption events compared to SED. That the SED gradient, a measure of fluid flow through the LCN, can improve predictions of remodeling events compared to SED, a measure of direct cell strain, has been suggested previously [16]. Furthermore, as the SED gradient encompasses the neighboring SED voxels, it provides information of a broader mechanical environment, which could explain the higher modeling performance observed with the SED gradient compared to SED. An additional limitation of this study was that the micro-FE analysis did not take into account the component of frequency. Although our approach enabled us to link bone remodeling events to mechanical environments *in vivo* at the local level, the addition of theoretical models that incorporate cellular mechanosensing and intercellular communication [12, 35–37] will be highly useful to improve our understanding of the relationship between loading frequency and bone adaptation across multiple scales.

In conclusion, these results suggest that bone adaptation is regulated by mechanical signals in the local *in vivo* environment and furthermore, that mechano-regulation is logarithmically dependent on loading frequency with frequencies below a certain threshold having catabolic effects, and those above anabolic effects. This study thereby provides valuable insights towards a better understanding of the mechanical signals influencing bone formation and resorption in the local *in vivo* environment.

## Materials and Methods

### Study Design

To investigate the effect of loading frequency on mouse caudal vertebrae, 11-week old female C57BL/6J mice were purchased (Charles River Laboratories, France) and housed at the ETH Phenomics Center (12h:12h light-dark cycle, maintenance feed and water ad libitum, three to five animals/cage) for one week. To enable mechanical loading of the 6^th^ caudal vertebrae (CV6), stainless steel pins (Fine Science Tools, Heidelberg, Germany) were inserted into the fifth and seventh caudal vertebrae of all mice at 12 weeks of age. After three weeks of recovery, the mice received either sham (0N), 8N static or 8N cyclic loading with frequencies of 2Hz, 5Hz, or 10Hz and were scanned weekly using *in vivo* micro-CT. All procedures were performed under isoflurane anaesthesia (induction/maintenance: 5%/1-2% isoflurane/oxygen). All mouse experiments described in the present study were carried out in strict accordance with the recommendations and regulations in the Animal Welfare Ordinance (TSchV 455.1) of the Swiss Federal Food Safety and Veterinary Office (license number 262/2016).

### Mechanical loading

The loading regime was performed for five minutes, three times per week over 4 weeks as described previously [14]. For the cyclic loading groups, sinusoidally varying forces (8N amplitude) were applied at 2Hz, 5Hz or 10Hz resulting in cycle numbers of 600, 1500 and 3000, respectively. For the static loading group, the force was maintained at 8N during the five minutes. For the sham-loaded group, the tails were fixed in the loading device for five minutes, but no loading was applied (0N).

### Micro-CT imaging and analysis

*In vivo* micro-CT (vivaCT 40, Scanco Medical AG, isotropic nominal resolution: 10.5 μm; 55 kVp, 145 μA, 350 ms integration time, 500 projections per 180°, scan duration ca. 15 min, radiation dose per scan ca. 640 mGy) images of the CV6 were acquired every week. Micro-CT data was processed and standard bone microstructural parameters were calculated in trabecular, cortical and whole bone by using automatically selected masks for these regions as described previously [26]. To calculate dynamic morphometric parameters, micro-CT images from consecutive time-points were registered onto one another. The voxels present only at the initial time point were considered resorbed whereas voxels present only at the later time point were considered formed. Voxels that were present at both time points were considered as quiescent bone. By overlaying the images, morphometrical analysis of bone formation and resorption sites within the trabecular region allowed calculations of bone formation rate (BFR), bone resorption rate (BRR), mineral apposition rate (MAR), mineral resorption rate (MRR), mineralizing surface (MS) and eroded surface (ES) [18].

### Micro-finite element (micro-FE) analysis

For each mouse at each time point, segmented image data was converted to 3D micro-FE models, with additional voxels added to the proximal and distal ends of the vertebrae mimicking intervertebral discs. All voxels were converted to 8 node hexahedral elements and assigned a Young’s modulus of 14.8 GPa and a Poisson’s ratio of 0.3 [14]. The bone was assumed to have linear elastic behaviour, which allowed for static loading in the micro-FE analysis [38]. The top was displaced by 1% of the length in z-direction (longitudinal axis), while the bottom was constrained in all directions. The micro-FE model was solved using a micro-FE solver (ParOSol). The results were then rescaled to an applied force of 8N for the loaded groups and 4N (physiological loading) for the sham-loaded group (0N) as described previously [39].

### Mechanical environment

The mechanical stimuli, which are hypothesized to drive load induced bone adaptation are deformation (direct cell strain) and interstitial fluid flow (shear stress) [27]. As a measure of the mechanical deformation, strain energy density (SED) magnitudes, defined as the increase in energy associated with the tissue deformation per unit volume, were analysed on the bone surface on the marrow-bone interface. Furthermore, based on the assumption that spatial differences in tissue deformation induce fluid flow, the spatial gradient of the SED was analyzed on the marrow side of the marrow-bone interface [28]. The spatial gradients in x, y and z-direction were calculated as follows:

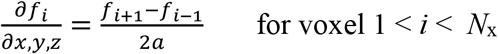

Where *f*_*i*_ is the SED of a voxel at x, y, z-position *i*, *N*_*x,y,z*_ the number of voxels in the x,y,z-direction and *a* the nominal resolution. The norm of the gradient vector (∇SED) was used as a quantity for the fluid flow as described previously [16].

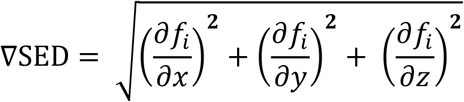

The conditional probabilities for a certain remodeling event (formation, quiescence, resorption) to occur at a given value of SED and ∇SED were calculated as described previously [15]. Briefly, the surface SED and ∇SED values were normalized within each animal and measurement by the maximal SED or ∇SED, respectively (chosen as the 99^th^ percentile of the values present at the surface and in the volume of interest (VOI) in order to remove the variance due to temporal bone adaptation, applied force in FE analysis and individual animals. For each region (formation, quiescence and resorption), a frequency density histogram with 50 bins and equal bin width was created. In order to rule out the dependence on the imbalance between bone formation and resorption, all remodeling events were assumed to have the same occurrence probability (i.e., formation, resorption and quiescent regions were rescaled to have the same amount of voxels). The remodeling probabilities were fitted by exponential functions using non-linear regression analysis.

To quantify the modeling performance of SED and ∇SED, respectively, the area under the curve (AUC) of a receiver operating characteristic (ROC) curve was used. The AUC can be defined as the probability that a randomly selected case (“true”) will have a higher test result than a randomly selected control (“false”) [40]. The ROC curve is a binary classifier, therefore the three different surface regions were analysed separately and only voxels and mechanical quantity values on the bone or marrow surface were used for the classification.

### Statistical analysis

Data are represented as mean±SD. For analysis of the longitudinal measurements of bone structural parameters, repeated measurements ANOVA implemented as a linear mixed model was used using the lmerTEST package [41] in R (R Core Team (2019), R Foundation for Statistical Computing, Vienna, Austria). The between subjects effect was allocated to the different groups (sham, static, 2Hz, 5Hz, 10Hz) while the within-subjects effects were allocated to time and time-group interactions. Random effects were allocated to the animal to account for the natural differences in bone morphometry in different mice. In cases where a significant interaction effect (group*time) was found, a Tukey post-hoc multiple comparisons test was performed. For comparisons between groups one-way ANOVA analysis followed by Tukey’s or Dunnet’s multiple comparisons test were performed as stated in the corresponding figure legends using SPSS (IBM Corp. Released 2016. IBM SPSS Statistics for Windows, Version 24.0. Armonk, NY, USA). The plots were created using GraphPad Software (GraphPad Prism version 8.2.0 for Windows, GraphPad Software, La Jolla California, USA). Significance was set at α<0.05 in all experiments.

## Acknowledgments

This manuscript is based upon work supported by the European Cooperation in Science and Technology (COST Action BM1402: MouseAGE), the European Research Council (ERC Advanced MechAGE ERC-2016-ADG-741883) and the Kyoto University Global Frontier Project for Young Professionals: the John Mung Program.

## Supplementary Information

**S1 Table. Summary of non-linear regression functions and corresponding coefficients for the conditional probability (SED and SED gradient) in trabecular bone for the different groups averaged over all time points.**

