## Supplementary information for "Mechano-regulation of bone adaptation is controlled by the local *in vivo* environment and logarithmically dependent on loading frequency"

**S1** **Table. Summary of non-linear regression functions and corresponding coefficients for the conditional probability (SED and SED gradient) in trabecular bone for the different groups averaged over all time points.**

|  | | Formation | | Resorption | |
| --- | --- | --- | --- | --- | --- |
| Regression line | | F = a*(1-exp(-b*SED/SEDmax))+y_0_ | | R = a*exp(-b*SED/SEDmax)+y_0_ | |
|  | | **SED** | **SED gradient** | **SED** | **SED gradient** |
| sham | a | 0.139 | 0.240 | 0.201 | 0.191 |
|  | b | 3.471 | 3.287 | 10.609 | 12.289 |
|  | y_0_ | 0.281 | 0.248 | 0.264 | 0.272 |
|  | R^2^ | 0.979 | 0.993 | 0.964 | 0.975 |
| static | a | 0.141 | 0.249 | 0.206 | 0.202 |
|  | b | 3.742 | 3.244 | 10.806 | 12.919 |
|  | y_0_ | 0.274 | 0.240 | 0.269 | 0.275 |
|  | R^2^ | 0.973 | 0.990 | 0.980 | 0.981 |
| 2Hz | a | 0.197 | 0.307 | 0.262 | 0.245 |
|  | b | 4.160 | 3.775 | 10.853 | 13.431 |
|  | y_0_ | 0.244 | 0.205 | 0.253 | 0.267 |
|  | R^2^ | 0.992 | 0.995 | 0.982 | 0.983 |
| 5Hz | a | 0.189 | 0.311 | 0.284 | 0.264 |
|  | b | 5.664 | 4.045 | 15.033 | 16.915 |
|  | y_0_ | 0.234 | 0.200 | 0.264 | 0.273 |
|  | R^2^ | 0.979 | 0.993 | 0.949 | 0.964 |
| 10Hz | a | 0.217 | 0.316 | 0.284 | 0.252 |
|  | b | 3.381 | 3.147 | 8.764 | 11.770 |
|  | y_0_ | 0.247 | 0.216 | 0.232 | 0.258 |
|  | R^2^ | 0.988 | 0.996 | 0.981 | 0.984 |
